# Proteomic analysis of Dhh1 complexes reveals a role for Hsp40 chaperone Ydj1 in yeast P-body assembly

**DOI:** 10.1101/016402

**Authors:** Gregory A. Cary, Dani B.N. Vinh, Patrick May, Rolf Kuestner, Aimee M. Dudley

**Affiliations:** Institute for Systems Biology, Seattle WA; Molecular and Cellular Biology Program, University of Washington, Seattle WA; Luxembourg Centre for Systems Biomedicine, Luxembourg; Pacific Northwest Diabetes Research Institute, Seattle, WA; Equally contributing author

## Abstract

P-bodies (PB) are ribonucleoprotein (RNP) complexes that aggregate into cytoplasmic foci when cells are exposed to stress. While the conserved mRNA decay and translational repression machineries are known components of PB, how and why cells assemble RNP complexes into large foci remain unclear. Using mass spectrometry to analyze proteins immunoisolated with the core PB protein Dhh1, we show that a considerable number of proteins contain low-complexity (LC) sequences, similar to proteins highly represented in mammalian RNP granules. We also show that the Hsp40 chaperone Ydj1, which contains an LC domain and controls prion protein aggregation, is required for the formation of Dhh1-GFP foci upon glucose depletion. New classes of proteins that reproducibly co-enrich with Dhh1-GFP during PB induction include proteins involved in nucleotide or amino acid metabolism, glycolysis, tRNA aminoacylation, and protein folding. Many of these proteins have been shown to form foci in response to other stresses. Finally, analysis of RNA associated with Dhh1-GFP shows enrichment of mRNA encoding the PB protein Pat1 and catalytic RNAs along with their associated mitochondrial RNA-binding proteins, suggesting an active role for RNA in PB function. Thus, global characterization of PB composition has uncovered proteins and RNA that are important for PB assembly.

## INTRODUCTION

Processing bodies (P-bodies, PB) and stress granules (SG) are eukaryotically conserved RNP granules consisting of non-translating mRNA and associated proteins (Eulalio, Behm-Ansmant, and Izaurralde 2007; Kedersha and Anderson 2009; Decker and Parker 2012). PB and SG both accumulate in cytoplasmic foci within minutes of exposure to a variety of environmental stresses (Teixeira *et al.* 2005; Brengues *et al.* 2005; Garmendia-Torres *et al.* 2014), and the appearance of these foci is correlated with global translational arrest common to the early phase of many cellular stress responses (Holcik and Sonenberg 2005; Kedersha and Anderson 2009; Simpson and Ashe 2012). The kinetics of assembly and exact composition of these granules can vary in a stress-specific manner (Buchan *et al.* 2011). PB and SG are primarily distinguished based on their constituent proteins; PB core proteins are associated with mRNA decay functions (Sheth and Parker 2003; Buchan *et al.* 2010), whereas SG consist of translation initiation factors as well as other mRNA binding proteins (Hoyle *et al.* 2007; Buchan *et al.* 2008; Lui *et al.* 2010; Decker and Parker 2012; Kedersha *et al.* 2013). These granules have been observed to interact *in vivo*, and mRNP sub-complexes can exchange between foci (Kedersha *et al.* 2005; Stoecklin and Kedersha 2013). Furthermore, specific proteins can cycle into foci from the cytoplasm in less than a minute (Aizer *et al.* 2008). These observations highlight the dynamic nature of RNP granule assemblies.

Regions of low-complexity (LC) sequence are common among proteins that localize to mammalian RNP granules (Kato *et al.* 2012). LC regions are necessary for both RNP aggregation into cytoplasmic foci (Kato *et al.* 2012) and retention of RNA (Han *et al.* 2012). Recent studies have also shown the prevalence of LC-domains in proteins that affect SG assembly in yeast (Yang *et al.* 2014). Similar Q/N-rich prion-like domains, a specific subset of LC domains, are found in several yeast PB and SG proteins (e.g. Lsm4, Edc3, and Pbp1) and are required for RNP granule aggregation (Gilks *et al.* 2004; Decker *et al.* 2007; Buchan and Parker 2009). Aberrant forms of these RNP granule proteins, with the propensity to form cytotoxic, prion-like aggregates, have been associated with a number of intractable degenerative diseases (Li *et al.* 2013; Ramaswami *et al.* 2013). While aggregation via LC sequences is a common feature for many RNP proteins, it is the result of a controlled physiological process as opposed to nonspecific protein-protein aggregation. For example, while salt stress induces PB aggregation in wild-type yeast, deletion of the gene encoding the effector kinase for the osmotic shock signal transduction pathway prevents the accumulation of PB foci in the presence of high salt (Teixeira *et al.* 2005).

To date, the functional relevance of RNP granule aggregation remains unclear. The fact that mutant strains that are unable to form foci show a decrease in cell viability (Lavut and Raveh 2012) and long term survival (Ramachandran *et al.* 2011) suggests that PB/SG aggregation has some important cellular function. Furthermore, PB foci can be transmitted from mother to daughter cells during yeast mitosis, and this transmission provides a measurable growth advantage to the recipient cells (Garmendia-Torres *et al.* 2014). However, despite the fact that PB consist of proteins involved in mRNA decay, mRNA decay processes are not affected by perturbations that block the formation of visible PB foci (Eulalio, Behm-Ansmant, Schweizer, *et al.* 2007; Decker *et al.* 2007). Thus, an inventory of the proteins and RNA transcripts that localize to these granules and a better understanding of how their composition changes in response to stress induction could shed light on the nature of the cellular benefit of PB/SG aggregation.

Much of our current understanding of the composition and assembly dynamics of PB and SG is based on cytological and genetic experiments that have characterized protein localization to cytoplasmic foci under stress conditions. Previous biochemical studies have identified few core proteins when purifying PB components under native conditions (Fenger-Gron *et al.* 2005; Drummond *et al.* 2011; Bahassou-Benamri *et al.* 2013), likely due to the dynamic nature of RNP granules. Other approaches to characterize PB components have relied on cross-linking and denaturing conditions to capture proteins and associated RNA (Mitchell *et al.* 2013). To better understand both the protein and RNA constituents of RNP granule complexes, we adapted a method to enrich native RNP complexes from yeast cells (Oeffinger *et al.* 2007) and used an anti-GFP antibody to isolate the complex associated with a GFP fusion to the core PB component, Dhh1. In order to characterize the components that differentially associate with PB during a stress and non-stress conditions, we isolated the Dhh1-GFP complex from cells grown in normal and acute glucose stress conditions and used quantitative tandem mass spectrometry (MS) and microarray analysis to identify the proteins and RNAs within the Dhh1 complex. Our results give evidence for PB association of many proteins previously implicated by genetic and cytological studies, and provide a new approach for analyzing the composition and function of these structures upon stress induction.

## MATERIALS AND METHODS

### Yeast strains and growth conditions

All yeast strains used in this study (Table S1) are derived from BY4741 (Winston *et al.* 1995). GFP-tagged strains (Huh *et al.* 2003) were purchased from Life Technologies. Individual gene deletions marked by *kanMX* were created by homologous recombination in strains harboring GFP-tagged genes (e.g. *DHH1-GFP*, YAD49). MoBY plasmids are from a library collection of CEN-copy plasmids that contain individual barcoded yeast ORFs expressed from their own promoter (Ho *et al.* 2009) that was purchased from ThermoScientific. Different MoBY plasmids were transformed into YAD557 and tested for Dhh1-GFP foci formation. All genetic manipulations of yeast and growth media are as in standard protocols (Rose *et al.* 1990). Unless otherwise noted, yeast cells were grown in rich media (YPD) at 30 °C.

For glucose depletion (-glucose) experiments, overnight 5 ml cultures in YPD were serially expanded into a final 2 L culture and grown to late log phase (OD_600_=1.0). Cells were rapidly harvested by filtration (Millipore Nitrocellulose Membrane; 0.65 µm pore size), and resuspended into fresh 2 L -glucose media (YEP), and the culture was shaken for an additional 30 min. Final cell pellets were collected by filtration, concentrated into conical tubes by low speed (500 x*g*) centrifugation, and flash-frozen in liquid nitrogen.

A SILAC culture of wild-type cells was generated using a modified I-DIRT (Tackett *et al.* 2005) protocol. Briefly, BY4741 was first grown to a late log density of OD_600_=1 in synthetic complete media lacking lysine and arginine (SC–lys–arg), containing 2% glucose, and supplemented with 50 mg/L each of lysine and arginine. This culture was then diluted and grown for a total of 9 doublings to OD_600_ ~1.2 in SC–lys–arg 2% glucose media supplemented with 50 mg/L each of heavy-isotopically labeled arginine (^13^C6-^15^N4) and lysine (^13^C6-^15^N2). Cells were filtered and frozen as described above.

### Cell lysis and immuno-affinity purification

Cell pellets stored at -80 ºC were released into a pre-cooled Retsch PM-100 planetary ball mill grinding jar. Grinding was performed at 600 rpm in 2-minute cycles with rotation reversals at 1 minute. Jars were re-chilled in liquid nitrogen between grinding cycles. Samples were ground until >90% lysis was achieved, which typically occurred after 5 to 10 cycles of grinding. The cell powder grindate was collected and returned to -80 ºC for storage.

We used anti-GFP IgG (Roche) coupled to magnetic Protein-G Dynabeads (Invitrogen) to capture Dhh1-GFP protein complexes from yeast cell lysate. Anti-GFP was first crosslinked to protein-G bead in 20 mM dimethyl pimelimidate (DMP; ThermoScientific) using protocols recommended by the manufacturer. A ratio of 30 mg of Ab-proG beads to 300 mg of total protein captured >90% of Dhh1-GFP in the cell lysate as determined by Western blot analysis. To obtain sufficient material for mass spectrometry, we used cell lysate generated from 4 L of cell culture. To generate a lysate supernatant, cell powder grindate was quickly thawed by resuspending in 1.5 volumes of RB buffer [30 mM K-Hepes (pH 7.4), 150 mM KCl, 2 mM MgCl_2_, 0.2% NP-40, 0.1% Tween-20, 1 mM DTT, yeast protease inhibitor cocktail (Sigma), and RNase-inhibitor (Ambion / Millipore)]. The suspension was subsequently cleared by low speed centrifugation (3,000 x*g*) for 5-7 min at 4 ºC. The resulting supernatant was incubated with anti-GFP-protG beads for 30 min at 4 ºC with gentle mixing. The beads were subsequently separated from the supernatant and washed extensively with RB buffer containing progressively less detergents [0.1% NP-40/0.05% Tween-20]. After the final wash, the beads were divided into separate fractions for elution of protein (88% of total) and RNA (12% of total). Protein was eluted in a solution of [0.1% SDS, 30 mM Hepes (pH 7.4), protease & RNase inhibitors] for 30 min at RT. A fraction of the eluted proteins was analyzed immediately by Silver staining (Pierce/ThermoScientific) and Western blot (Licor Odyssey) using a different anti-GFP antibody (Clontech), while the remaining was flash frozen in liquid nitrogen and stored at -80 °C. The identities of some of the dominant bands on silver stained SDS gels (Figure 1, marked with stars) are PB components assigned by their apparent molecular weight by gel migration and mass spectrometry (MS) from other purifications. For RNA isolation, beads were incubated with [1% SDS, 30 mM Hepes (pH 7.4), protease & RNase inhibitors] for 30 min at RT. The resulting supernatant was mixed with an equal volume of TES buffer [1% SDS in 10 mM Tris (pH 7.5), 1 mM EDTA], added to a total equal volume of acid phenol (pH 4), and incubated at 65 °C for 60 min. Samples were spun at 15K x*g,* 5’, then the aqueous phase was collected and extracted again with phenol:chloroform, followed by precipitation with cold ethanol. Final pellets were resuspended in TE and stored at -80 °C.

**Figure 1.**
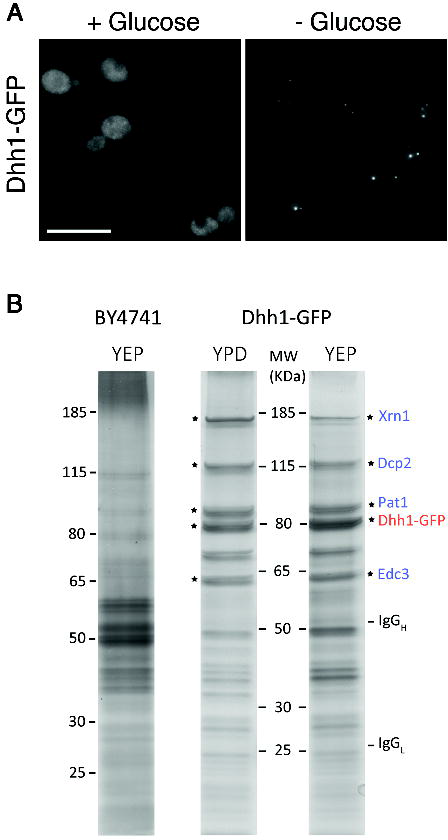
Immunoisolation of Dhh1-GFP. (A) Fluorescence microscopy showing the aggregation of Dhh1-GFP into cytoplasmic foci in cells grown in media with and without glucose for 30’. Scale bar = 10 µm. (B) Silver stained SDS-PAGE gels of IP fractions of Dhh1-GFP isolated from cells grown in +glucose (YPD) and -glucose (YEP) media; and from the negative control BY4741 cells grown in -glucose. The identities of some of the dominant bands (marked with an asterisk, Dhh1-GFP (red), and dominant co-isolated PB proteins (blue)) are based on the apparent molecular weight and by mass spectrometry from other purifications. MW: molecular weight of standard proteins.

For I-DIRT experiments, isotopically-heavy BY4741 and light Dhh1-GFP frozen cell pellets were ground separately and equal weights of each grindate (to generate 1:1 total protein mass) were mixed immediately before resuspending in RB buffer. The suspension was then cleared by centrifugation and the supernatant collected. Immuno-isolation with the anti-GFP antibody was performed as described above.

### Mass spectrometry and proteomic analyses

Eluted fractions from the anti-GFP immunoprecipitation were first TCA precipitated, then reduced, alkylated, and trypsinized (Promega) (Tian *et al.* 2007). Tryptic digestions were acidified and then desalted by UltraMicroSpin Vydac C18 silica column (Nest Group, Inc.) following manufacturer’s specifications. Desalted samples were dried and resuspended in a solution of 5% acetonitrile and 0.1% formic acid prior to tandem mass spectrometric analysis on an LTQ-Velos (for spectral counts) or LTQ-Orbitrap (for I-DIRT) electrospray ionization mass spectrometer.

Tandem mass spectra were converted to universal mzXML file format and searched against a database consisting of all known yeast open reading frames (Saccharomyces Genome Database; SGD) (Costanzo *et al.* 2014), known contaminant proteins, and a decoy library prepared by randomization of the library using a perl script available from the Matrix Science website (http://www.matrixscience.com/help/decoy_help.html). Searches were performed using the program X!Tandem (Craig and Beavis 2004) with the following parameters: tolerable tryptic termini = 1; identifications based on b- and y-ions; parent mass tolerance = 3.00; fixed modifications include carboxyamidomethylation of cysteine (MW= 57.02); variable modifications include oxidation of methionine (MW= 15.99). Tandem mass spectra peptide and protein assignments were validated by the PeptideProphet (Keller *et al.* 2002) and ProteinProphet (44) programs, available in the current TPP distribution (http://tools.proteomecenter.org/wiki/index.php?title=Software:TPP). Protein probabilities yielding a 0.05 FDR threshold were applied to the resulting protein lists and filtered to exclude proteins identified in any experiment with fewer than two unique peptides.

For spectral count quantitation experiments, paired -glucose and +glucose immunopurified samples were analyzed with two technical replicate injections for each sample on the same mass spectrometer on the same day. Because the number of measured spectra can vary depending on criteria such as protein length or primary sequence, tandem mass spectral counts for each protein identified were normalized using the APEX program (45, 46) (version 1.1.0) available from the JCVI website (http://pfgrc.jcvi.org/index.php/bioinformatics/apex.html). Pre-calculated observability scores for the yeast proteome (yeast_ORBI_66attrib_ALLpredictions.Oi) were downloaded (http://marcottelab.org/APEX_Protocol/Oi_Predictions/Scerevisiae) and input along with prot.xml files output by the TPP. All proteins lower than the 0.05 FDR threshold were APEX-normalized and pairs of -glucose and +glucose samples were compared using the two-sample Z test utility. The APEX-normalized value for each protein was further normalized to the APEX score of Dhh1 from the same sample.

The mass spectrometric data from the I-DIRT immunoprecipitated samples were analyzed using a similar approach as described above with the following differences. Database searches were performed using the following variable modifications: SILAC heavy arginine: ^13^C6-^15^N4 (MW= 10.01), and SILAC heavy lysine: ^13^C6-^15^N2 (MW= 8.01). Following assessment of peptide and protein identifications by PeptideProphet (Keller *et al.* 2002) and ProteinProphet (Nesvizhskii *et al.* 2003), quantitative SILAC ratios for proteins were determined using XPRESS software (Han *et al.* 2001). Precursor ion elution profiles of heavy versus light peptides were determined with a mass tolerance of 0.05 (>5s) and the area under the curve was used to determine a SILAC ratio for each peptide.

Two criteria were used to filter the results down to the final list of 329 proteins. First, only proteins identified in any experiment with more than two unique peptides were considered; and second, proteins identified in at least any two of the five + and -glucose experiments (313 proteins), or identified in any one of the five experiments plus measured in at least one of the two I-DIRT replicates with a light:heavy SILAC ratio greater than 50:50 (an additional 16 proteins) were included. Gene ontology (GO) category enrichments of the isolated proteins were determined using the YeastMine toolset available from the SGD website (http://yeastmine.yeastgenome.org/yeastmine/). All reported *p*-values were corrected for multiple hypothesis testing using a Benjamini-Hochberg correction (Benjamini and Hochberg 1995).

### Microarray analysis of RNA enrichment

A custom Agilent DNA microarray was designed that consists of 30,529 probes antisense to the *S. cerevisiae* transcriptome (Agilent Design ID: 045101). The probes on the array were designed to hybridize to 10,283 different yeast transcripts including the 6,607 ORF transcripts annotated in the Saccharomyces Genome Database as well as 3,676 non-coding RNAs. For each transcript, three distinct 60 bp probes were designed and distributed across the array. In total, seven experiments were assessed by microarray analysis, including RNA extracted from all five preps used to generate protein for MS analysis (above), as well as one additional -glucose sample and a mock IP of lysate prepared from a strain expressing GFP alone. For each experiment, a sample of RNA (total RNA) was extracted from cell lysate before immuno-purification and compared to RNA extracted from the immuno-purified complexes (IP RNA). 5 µg of total RNA and 200 ng IP RNA were fragmented and hybridized per manufacturer’s directions to two separate microarrays for each pair of total and IP RNA samples. We used an antibody-based method to directly detect RNA:DNA hybrid on the array (Dutrow *et al.* 2008). The primary S9.6 antibody was purified from hybridoma cell line (ATCC clone HB- 8730). Secondary antibody detection was with a Cy3-labeled anti-mouse antibody. Following the final antibody wash, slides were dried by brief low speed (600 rpm) centrifugation and immediately scanned. Feature extraction was performed using an Agilent G2565CA Microarray Scanner and control software. The median background signal from 1,559 array features designed to have no homology to yeast transcripts was subtracted from all features. Background subtracted signal was log transformed, pairs of Total and IP arrays were normalized by cyclic loess implemented in the Limma Bioconductor package (Smyth 2005), and transcript replicate probes were averaged. Finally, we applied a threshold to remove transcripts that exhibited low abundance signal and high variability or that were saturated on the total RNA array. We generated a linear regression between the average normalized transcript signal in each IP RNA sample and the signal from that transcript in the matched total RNA sample and considered a transcript enriched by the IP if it lay outside the upper 95% confidence interval of the regression line. Microarray design information and data have been deposited on the Gene Expression Omnibus (GEO) database at NCBI (Accession number: GSE65989).

### Microscopy and image analysis

Cells were grown to mid-log phase (OD_600_ ~ 0.7) in YPD, then pelleted, washed in YEP media without glucose, resuspended in YEP, and grown for another 30 minutes. Cells were fixed in 2% para-formaldehyde (MeOH free; Polysciences), 10 minutes at room temperature. Cells were then washed and stored in 1.2M sorbitol/0.1M K-phosphate (pH 7.5). Fixed cells were imaged using a DeltaVision microscope system (Applied Precision, Issaquah, WA), through a 60x oil objective lens in the Olympus IX-71 wide field microscope. Sets of 30, 0.2 micron z-sections were captured for each image, then deconvolved using softWoRx® software (Deltavision trademark). Finally, ImageJ software (imagej.nih.gov/ij) was used to adjust contrast levels and images in all the stacks collapsed into one final image.

## RESULTS

### Isolation of P-body components

Adapting a comprehensive approach to analyze the composition of intact RNP complexes (Oeffinger *et al.* 2007), we developed an immuno-affinity method to isolate the PB core component Dhh1 at high yield under conditions that best preserve PB integrity. We chose Dhh1-GFP because of its abundance relative to other components (Ghaemmaghami *et al.* 2003), it interacts with several PB components (Coller *et al.* 2001), and is a component of both yeast PB and SG (Buchan *et al.* 2008; Swisher and Parker 2010). Similar to previous studies, Dhh1-GFP appears cytoplasmically diffuse when cells are grown in media containing glucose but rapidly aggregates into cytoplasmic foci upon an acute stress of glucose depletion (Figure 1A). Cells grown in these two media were rapidly collected by vacuum filtration and flash frozen in liquid nitrogen to minimize induction of PB aggregation during the sample preparation. To best preserve PB sub-complexes and to avoid protein and RNA degradation, cells were lysed in this frozen state by planetary ball mill grinding in liquid nitrogen. Intact Dhh1-GFP foci were observed in the generated supernatant by light microscopy. Finally, high yields of Dhh1-GFP complexes (>95% depleted from supernatant) were isolated under native conditions using high affinity anti-GFP antibodies coupled to protein-G magnetic beads.

Silver-stained gels of eluted proteins indicate an enrichment of Dhh1-GFP and associated proteins that are not observed in the control BY4741 sample (Figure 1B). Several core components of the mRNA decay complex (Pat1, Edc3, Dcp2, and Xrn1) co-purified with Dhh1 based on their apparent molecular weights. The intensities of these major bands correlate well with their being some of the most abundant proteins in all Dhh1-GFP complexes as measured by mass spectral counts (discussed below).

### Proteomic analysis of Dhh1-GFP complexes

To compare PB composition between different cell growth conditions, we analyzed a total of five Dhh1-GFP purifications (two from cells grown in +glucose media, three from -glucose media) by mass spectrometry. In addition, two SILAC-based purification experiments, termed I-DIRT (Tackett *et al.* 2005), were conducted to further assess *in vivo* protein interactions with Dhh1-GFP. In this approach, co-immunoprecipitated proteins that were labeled with isotopically-heavy amino acids from control cells without GFP can only associate with Dhh1-GFP during light- and heavy-labeled extract mixing *in vitro* (Materials & Methods). As shown in Figure S1 and Table S2, most PB proteins show strong *in vivo* interactions with Dhh1-GFP, with light:heavy peptide ratios greater than 50:50 (log2 XPRESS ratio > 0, see Materials & Methods). Interestingly, some known PB components, including Dcp1 and Dcp2, appear to have neutral to low light:heavy peptide ratios, suggesting a dynamic exchange with the isolated Dhh1-GFP complexes, consistent with mammalian studies *in vivo* (Aizer *et al.* 2008). Therefore, we chose not to exclude proteins with light:heavy peptide ratios lower than 50:50 as these might include important components of the complex with dynamic exchange rates. Instead, we considered SILAC light:heavy peptide ratios higher than 50:50 as additional evidence supporting *in vivo* interactions.

In total, we identified 329 proteins that were statistically significant by mass spectrometric analyses and associated with Dhh1-GFP reproducibly in the biological replicates (Materials & Methods; Table S2). There are a number of proteins that overlap between conditions but are not identified in all replicates of similar condition (*i.e.* -glucose condition; Figure S2) suggesting that while the profile of major core proteins in Dhh1-GFP complexes appear similar on SDS-PAGE, the less abundant interacting proteins may vary between similar conditions. Therefore, only the 329 proteins that were identified in at least two biological replicates from any condition were considered for further analysis.

### Proteins known to associate with PB

Strikingly, about 16% of the total 329 Dhh1-interacting proteins have previously been linked to PB and SG. In many cases, our results provide biochemical evidence for proteins that have been associated with PB or SG only by co-localization or genetic studies. Among the enriched PB and SG proteins, 19 are considered core components (Table 1), (Buchan *et al.* 2010). Most known PB core components are identified, and relative to all proteins coisolated with Dhh1-GFP, the family of decapping proteins Dcp1, Dcp2, Edc3, Pat1, Lsm1-7, and exonuclease Xrn1, are the most abundant PB proteins whether isolated from cells grown in -glucose or +glucose media (highest spectral counts, Table S4). Similar to the SDS-PAGE profile (Figure 1), the stoichiometry among these 12 core proteins relative to each other and to Dhh1 appears very similar in both growth conditions, suggesting that an inherent Dhh1 core sub-complex exists regardless of the induction status of PB/SG foci. These results are consistent with previous studies showing that several core PB components including Dhh1 can bind directly to each other by *in vitro* pull-down assays or immunoprecipitations from cells grown in glucose supplemented media (Tharun *et al.* 2000; Coller *et al.* 2001; Kshirsagar and Parker 2004; Decker *et al.* 2007; Harigaya *et al.* 2010). Notably Lsm1, and not Lsm8, was identified along with the Lsm2-7 proteins in all of the purification replicates. Lsm1 binds to Lsm2-7 proteins and forms a cytoplasmic decapping complex that associates with PB, while Lsm8 forms another complex with Lsm2-7 that is recruited to the nucleus to perform splicing and processing of nucleolar and ribosomal RNA (Novotny *et al.* 2012). The absence of Lsm8 in our purifications suggests a preferential association of cytoplasmic Lsm1-7 complexes with Dhh1-GFP. Core SG proteins that interact with Dhh1-GFP (Table 1) are also identified, but at lower abundance (by spectral counts) than core PB proteins (Table S4). Finally, 34 coisolated proteins are identified as PB/SG-associated (Table 1) because they colocalize partially with PB/SG core proteins, or they affect PB/SG assembly (Balagopal and Parker 2009; Buchan and Parker 2009; Tkach *et al.* 2012; Buchan *et al.* 2013; Yang *et al.* 2014). Due to the dynamic interchange of components between PB and SG foci, we will denote Dhh1-GFP binding proteins hereon as PB/SG.

**Table 1.**
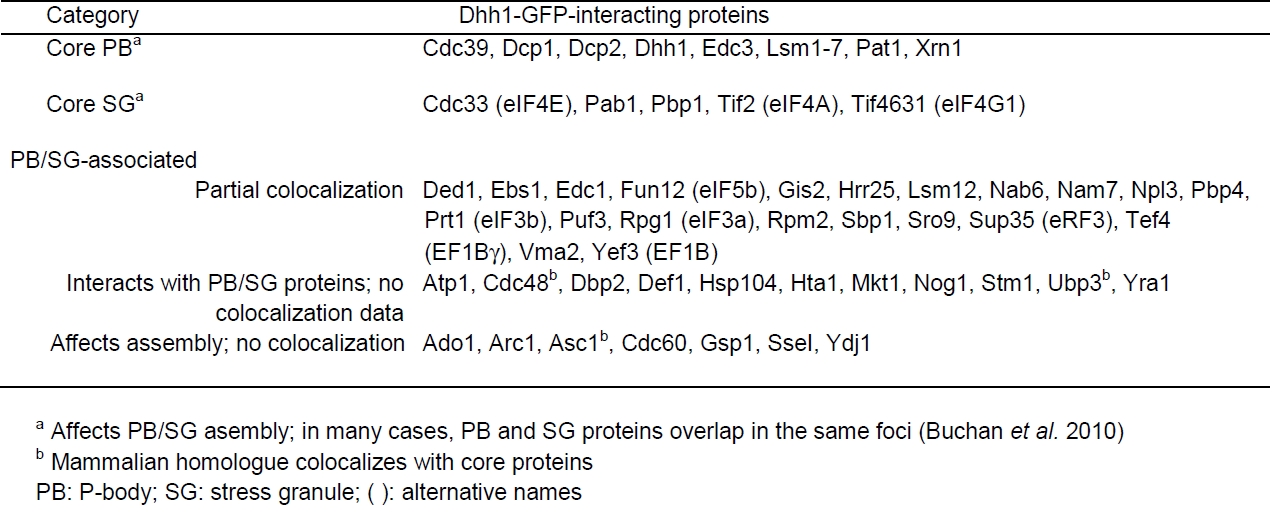
Known P-body and stress granule components co-isolated with Dhh1-GFP

One of the most enriched categories of cellular components from our list of 329 Dhh1 associated proteins was a set of ribosomal proteins (structural constituent of ribosome, 85 total; hypergeometric p-value=9x10^-42^). This result is consistent with several studies showing a close association between ribosomes and PB components, including Dhh1 (Kressler *et al.* 1997; Bonnerot *et al.* 2000; Hu *et al.* 2009; Drummond *et al.* 2011; Sweet *et al.* 2012; Cougot *et al.* 2012). Components of both the 40S and 60S subunits were identified, including Rps30A and Rpl16A (Table S2) which have been reported to bind Dhh1 (Drummond *et al.* 2011; Sweet *et al.* 2012). More than 40% of the Dhh1-GFP interacting proteins are non-ribosomal RBP that are part of ribonucleoprotein (RNP) granules (p=6x10^-18^) (PB and SG) and RNP complexes (p=6x10^-14^). These include other preribosomal, polysomal, nucleolar, and translation associated proteins (Figure 2A, Table S3). In addition to their known targets of rRNA, tRNA or snoRNA for ribosome biogenesis, or their role in other cellular functions such as vacuolar trafficking or glycolysis, many of the 120 non-ribosomal RBP have recently been found to bind mRNA (Hogan *et al.* 2008; Scherrer *et al.* 2010; Tsvetanova *et al.* 2010; Mitchell *et al.* 2013). Thus, affinity purifying Dhh1 under native condition has allowed us to isolate the core PB subcomplexes along with many PB/SG associated proteins known for their role in mRNA binding and translational functions.

**Figure 2.**
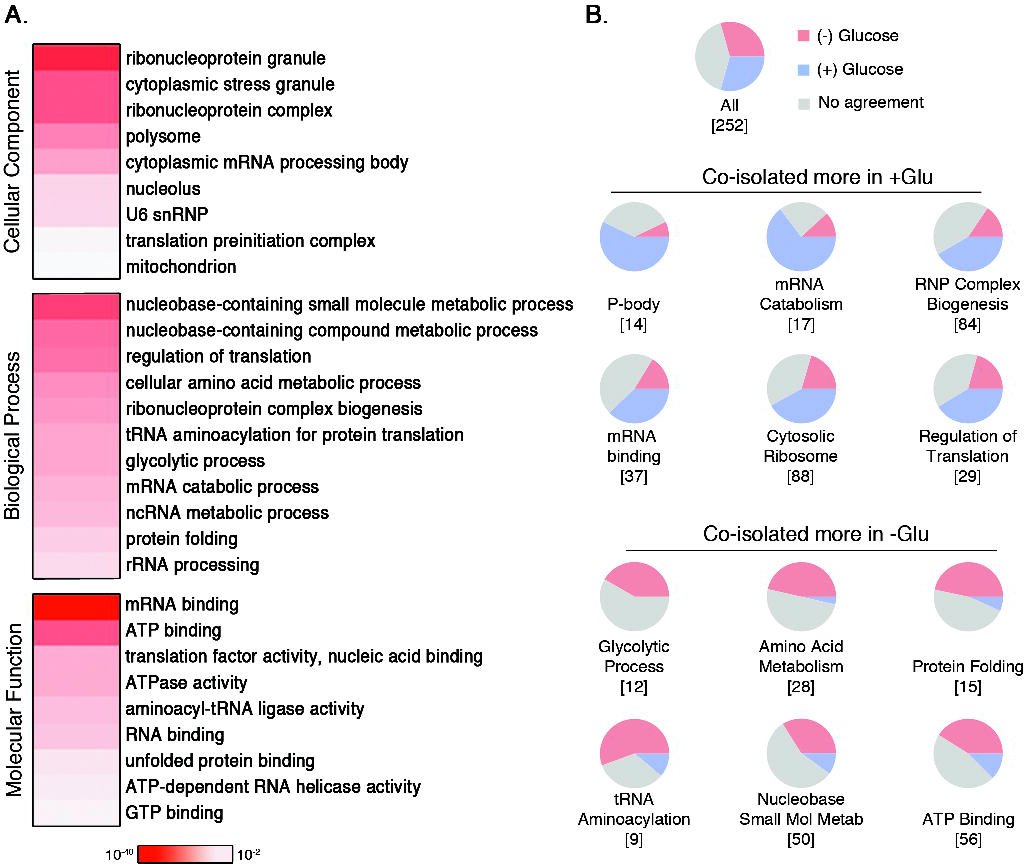
Protein constituents of Dhh1-GFP complexes. (A) Gene ontology (GO) enrichment represented as heatmap of 244 non-ribosomal Dhh1-GFP interacting proteins. Color intensity corresponds to the p-value from the hypergeometric test after correcting for multiple hypothesis testing. (B) Pie charts show the proportion of each GO category that reproducibly change in abundance when coisolated with Dhh1-GFP in different growth conditions. Protein abundance was determined using APEX-normalized spectral counts and further normalized to the level of Dhh1-GFP. Proteins that are reproducibly more abundant when coisolated with Dhh1-GFP from cells grown in +glucose are shown in blue, those reproducibly more abundant when coisolated from cells grown in -glucose are shown in red, or where there is no agreement between replicates in grey. The total number of proteins from each category measured in both spectral count experiments is indicated in square brackets.

### Metabolic enzymes, tRNA synthases and protein chaperones

In addition to known RNA granule, ribosomal, and RNA binding proteins, the 244 non-ribosomal proteins were enriched for several other functional classes. One class included proteins associated with metabolic and biosynthetic processes, in particular proteins annotated as binding ATP or GTP (e.g. “nucleobase-containing small molecule metabolic process”, p=4x10^-15^) (Figure 2A). About 40% of the proteins included in this latter GO are also RBP (Table S3). Other related GO processes include cellular amino acid metabolism (p=8x10^-10^), glycolysis (p=2x10^-8^), protein folding or chaperones (p=9x10^-6^), and tRNA aminoacylation (p=2x10^-8^). Because proteins in these classes have been shown to localize to cellular foci in response to stresses such as DNA replication inhibition, acute glucose depletion, and stationary phase starvation (Narayanaswamy *et al.* 2009; Tkach *et al.* 2012; Mitchell *et al.* 2013), we counted the number of our 329 Dhh1-associated proteins for which there was previous evidence of stress-dependent localization to cellular foci.

We found evidence in the literature that 62 of our 329 Dhh1-GFP associated had been observed to form foci in at least one of the three stress conditions (Table S5), including 30 of the known PB/SG components in our list. Of the remaining 32 Dhh1-GFP interactors that had not been shown to co-localize with PB/SG proteins, 22 proteins, including metabolic enzymes, tRNA synthetases, and protein chaperones, form foci only in stationary phase cells. Taken together, these findings suggest that a subset of proteins that respond to stress by re-localizing to foci interact with Dhh1-GFP and are thus new candidates for PB/SG components.

### Proteins that preferentially associate with Dhh1-GFP in the PB induced state

To compare the composition of the Dhh1 complex in the two different cell growth conditions, we analyzed pairs of Dhh1-GFP immuno-isolations from -glucose and +glucose media detected by mass spectrometry using the same instrument on the same day to mitigate background variability. We use the APEX program (Braisted *et al.* 2008; Vogel and Marcotte 2008) to determine the normalized spectral count for each protein as an indication of their relative abundance in the protein purification (see Materials & Methods). APEX spectral count scores for each protein were further normalized to the APEX spectral count of Dhh1-GFP itself measured in the same sample, and the ratio of normalized scores from pairwise -glucose to +glucose samples was generated as a measure of the extent to which each protein is preferentially associated with Dhh1-GFP in the -glucose condition (PB induced) (Table S4). Because we limited this analysis to only those proteins that were identified in both replicates of the pairwise experiments, only 252 out of our 329 total proteins were included in the analysis. Of these, 71 proteins had scores consistent with higher levels in the +glucose condition and 71 proteins had scores consistent with higher levels higher levels in the -glucose condition. The set of proteins predicted to be associated with Dhh1 more in the +glucose condition had significant enrichment for the GO categories of RNA catabolism, RNP biogenesis, and regulation of translation. In contrast, those predicted to associate at higher levels in the -glucose condition (PB induced) were significantly enriched for glycolysis, amino acid metabolism, protein folding (chaperones), tRNA animoacylation (tRNA synthetases), and ATP-binding (Figure 2B). These results suggest that the induced state of PB tends to accumulate similar classes of PB/SG components that respond to stress by re-localizing to foci.

### Enrichment of proteins with low-complexity and prion domains in yeast RNP granules

Studies in mammalian cells have identified low-complexity (LC) regions as a common feature of many RBP and demonstrated the importance of these regions for both RNP granule aggregation and RNA retention (Han *et al.* 2012; Kato *et al.* 2012; Castello *et al.* 2012). We tested whether our set of Dhh1-associated proteins contained LC domains by two orthogonal approaches. First, we searched the set of 5887 ORFs annotated in the yeast genome database for LC sequences of at least 35 residues using the SEG algorithm (Wootton 1994) and found that 390 proteins meet these criteria (Table S6). Similar to the findings in mammalian systems (Kato *et al.* 2012), these proteins are strongly enriched for mRNA binding functions (p=3x10^-10^) and regulation of gene expression (2x10^-13^). Second, we considered a set of 178 yeast proteins predicted by Alberti *et al*. (Alberti *et al.* 2009) that contain putative prion-like domains. Protein-protein interactions via prion-like Q/N-rich domains, a specialized class of LC domains, have been shown to be essential for PB and SG aggregation (Gilks *et al.* 2004; Decker *et al.* 2007; Reijns *et al.* 2008). Both lists (390 LC and 178 prion) share 72 proteins in common (p=1.6x10^-44^), suggesting that these approaches identify similar protein sequence features (Table S6).

Of the 390 proteins predicted to contain LC domains, 42 were co-isolated with Dhh1-GFP (p=4.8x10^-7^). Of the 178 proteins predicted to contain putative prion domains, 29 were co-isolated with Dhh1-GFP (p=2.5x10^-9^). In total, we found that 55 Dhh1-GFP interactors contained predicted LC or prion-like domains, including 16 proteins that were in both sets. Of these 55 proteins, 38 are known RBP and the most significantly enriched GO categories include mRNA binding, RNP complex, RNP granule, and RNP complex biogenesis (Table S6). These results are consistent with mammalian studies demonstrating that proteins containing LC regions are highly represented in yeast RNP granules such as PB and SG.

### Ydj1 regulates Dhh1-GFP foci formation under acute glucose depletion stress

We identified 19 protein chaperones that co-isolate with Dhh1-GFP, including components of the CCT/TRiC chaperonin complex and members of the Hsp40, Hsp70 and Hsp90 families. Members of these chaperone families have been shown to be involved in PB/SG assembly by regulating the interactions between Q/N domain proteins (Gilks *et al.* 2004; Matsumoto *et al.* 2011; Nadler-Holly *et al.* 2012). Given that a significant number of proteins co-isolating with Dhh1-GFP also contain putative LC/prion domains, we tested whether any of the chaperones identified might be involved in regulating the aggregation of PB/SG foci. Because protein chaperones as a group are associated with Dhh1-GFP at a higher level in cells grown in -glucose media (PB induced condition) (Figure 2B), we focused on this condition.

The Hsp40 family protein Ydj1 has an LC domain (Table S6), binds Q/N-prion domain proteins (Summers *et al.* 2009), and in our proteomic studies associates with Dhh1-GFP only in the glucose depleted condition (Table S4). To test the effect of Ydj1 on PB assembly, we examined the ability of Dhh1-GFP, Lsm1-GFP or Edc3-GFP to localize to cytoplasmic foci in a *ydj1Δ* mutant background. In contrast to wild-type cells, Dhh1-GFP or Lsm1-GFP foci do not form when *ydj1*Δ mutant cells are grown in glucose-depleted media (Figure 3A). The formation of Dhh1-GFP or Lsm1-GFP foci when cells were grown to saturation, another PB inducing condition, was also drastically reduced in the *ydj1*Δ mutant background. In contrast, the loss of *ydj1* had minimal effects on Edc3-GFP aggregation (Figure 3A). The Dhh1-GFP foci assembly defect in *ydj1*Δ mutant (YAD557) was complemented by plasmids containing a wild-type copy of *YDJ1*, but not any of the other protein chaperones isolated in our Dhh1-GFP purifications (Figure 3B and 3C). We also tested whether deletion of other members of the Hsp70, Hsp90 and Hsp110 families, including *ssa1*Δ, *ssa2*Δ, *hsc82*Δ, *hsp82*Δ, or *hsp104*Δ, affected the stress induction of Dhh1-GFP foci and found that Dhh1-GFP foci formation to be the same as wild-type in all these mutant backgrounds (Figure S3). Finally, overexpression of either *YDJ1* or *SIS1*, two members of Hsp40 family that co-purified with Dhh1-GFP, did not induce the aggregation of either Lsm1-GFP or Dhh1-GFP foci in the absence of a glucose depletion stress [data not shown], suggesting that Hsp40 chaperones do not induce aggregation of PB/SG foci in the absence of an additional cellular stress. Taken together, these results suggest that Ydj1 is required for the formation of Dhh1- and Lsm1-containing cytoplasmic foci in response to glucose limited stress conditions.

**Figure 3.**
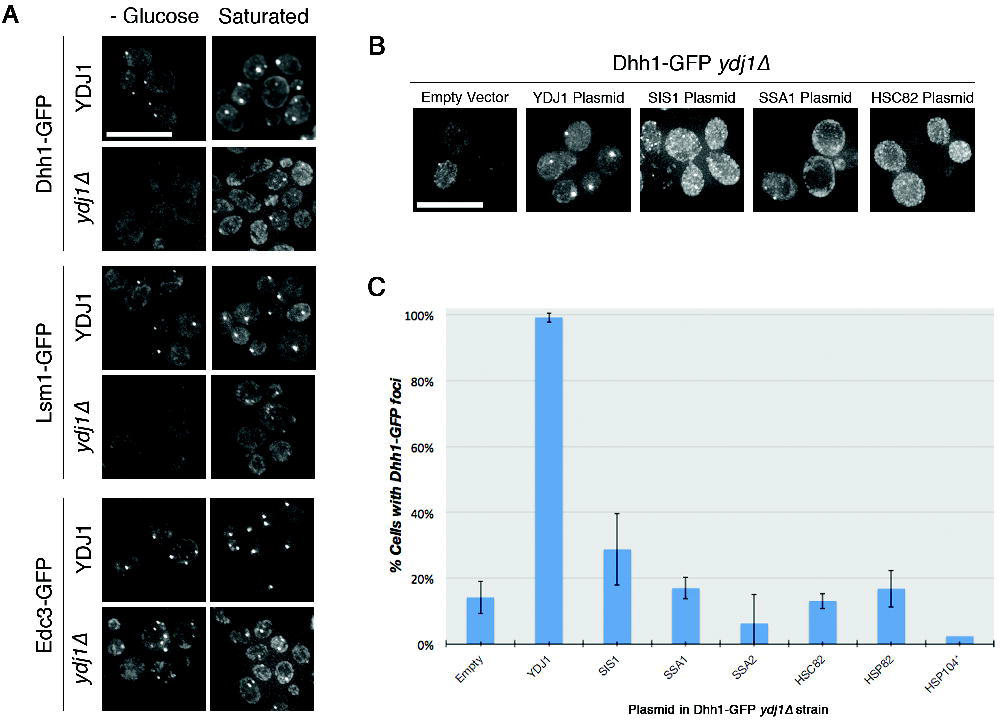
Ydj1 is necessary for the formation of PB foci. (A) Fluorescence microscopic images of Dhh1-GFP, Lsm1-GFP or Edc3-GFP in wild type (*YDJ1*) or mutant (*ydj1*Δ) cells following a 30-minute glucose depletion, or overnight culture. Scale bar = 10µm. (B) Microscopic images of Dhh1-GFP in *ydj1*Δ mutants transformed with CEN plasmids harboring the corresponding genes. All strains were induced to form foci by 30 minutes of glucose depletion. Scale bar = 10µm. (C) Quantitation of the percent of cells with at least 2 Dhh1-GFP foci following 30 minutes of glucose depletion for the strains in Figure 3B. Data presented are the average of at least two replicate experiments in which a minimum of 40 cells were counted. Error bars represent the standard deviation of the replicate measurements. (* = only 1 replicate quantified for Hsp104).

### Analysis of RNA associated with Dhh1-GFP complexes

Because PB are known to depend on the presence of RNA for their integrity (5), we isolated RNA in parallel from each Dhh1-GFP immuno-purified sample that was analyzed by mass spectrometry. To facilitate the analysis of strand-specific transcripts that were not dependent on the polyadenylation status of the transcript or biased by the amplification protocol, we hybridized RNA directly to a custom DNA microarray and measured RNA abundance using an antibody specific for RNA:DNA hybrids (Hu *et al.* 2006; Dutrow *et al.* 2008) (details in Materials & Methods). Transcripts were considered enriched in the IP if they were above the 95% confidence interval of a linear regression between the normalized signal from matched IP and total RNA samples (Figure 4A). Using this method, we identified 79 transcripts that specifically co-isolated with Dhh1-GFP in more than one IP biological replicate, but not in the mock sample which immuno-precipitated GFP alone (Table S7). The majority of these 79 transcripts are significantly enriched for non-coding RNAs, including dubious ORF transcripts, CUTs, SUTs (Xu *et al.* 2009), and MUTs (Lardenois *et al.* 2011) (Figure 4B). We also detect evidence of enrichment of Ty retroelements, along with proteins encoded by Ty elements, consistent with Ty association with PB (Checkley *et al.* 2010). While the majority of identified transcripts are noncoding, we did not detect co-enrichment of related transcripts, e.g. pairs of sense and antisense transcripts, which would suggest a regulatory role.

**Figure 4.**
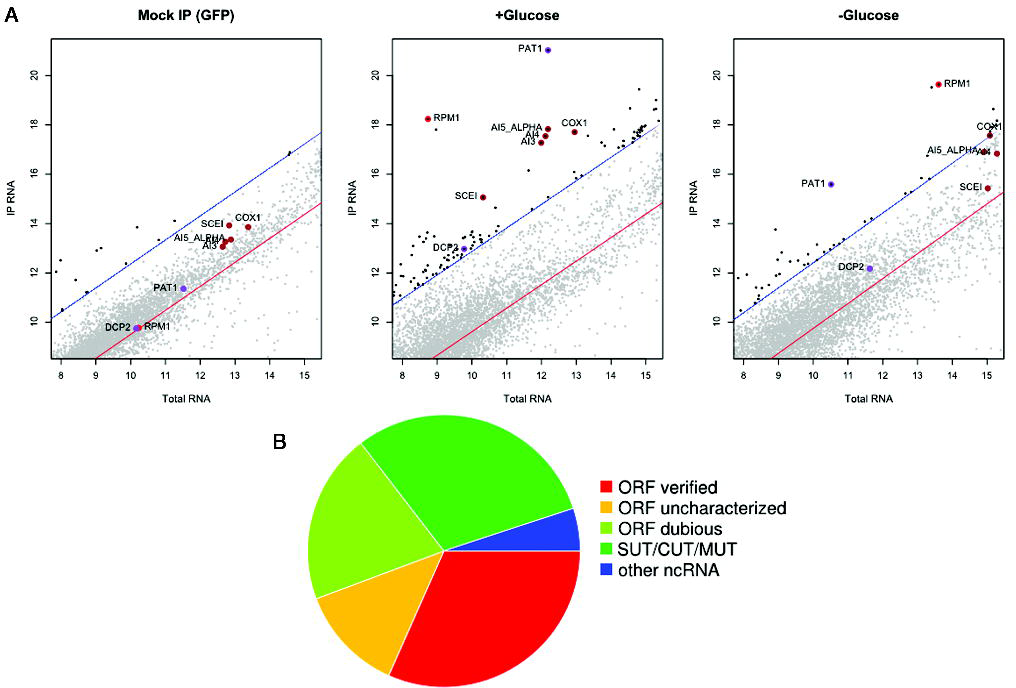
Analysis of RNA isolated in Dhh1-GFP complexes. (A) Representative microarray data from a mock IP (GFP alone) as well as Dhh1-GFP immunoisolations from a +glucose and a -glucose condition. The x-axis in each plot is the log-transformed, normalized array intensity for input (Total) RNA and the y-axis is the log-transformed, normalized array intensity for IP RNA. In each plot, the red line is a linear regression between the signals from the IP and total RNA, and the blue lines correspond to the 95% confidence interval. Transcripts above 95% confidence interval are considered enriched (and shaded black). Specific transcripts that are enriched and discussed in the text are highlighted and labeled (red= *RPM1*; brown= mitochondrial introns; purple= *PAT1* and *DCP2*). (B) The proportion of each RNA class within the total identified 79 transcripts. ORF= open reading frame; SUT= stable unannotated transcript; CUT= cryptic unannotated transcript; MUT= meiotic unannotated transcript.

Interestingly, the most enriched transcript in every Dhh1-GFP IP replicate is the mRNA encoding the PB protein Pat1 (Figure 4 and Figure S4B). mRNA encoding Dcp2 is also enriched, but not to the same extent as the PAT1 mRNA (Figure 4). Pat1 and Dcp2 are among the most abundant PB proteins in the Dhh1-GFP purification. Co-enrichment of both protein and mRNA could occur if nascent translation products remain associated with the mRNA on polysomes or if the protein regulates its cognate transcript (Pullmann *et al.* 2007). Of all the 329 Dhh1-GFP interacting proteins, only 9 mRNAs are found together with their cognate proteins in the same IP replicate (Figure S5), suggesting that transcripts are generally not isolated together with their cognate proteins in these immunoprecipitations. Pat1 is unique among the PB core proteins in that it can be shuttled between the nucleus to cytoplasm and this localization depends on associating with PB proteins Lsm1 and Dhh1 (Teixeira and Parker 2007; Hurto and Hopper 2011; Bahassou-Benamri *et al.* 2013), suggesting Pat1 has a nuclear function that may not be directly associated with PB. This would be consistent with the behavior of the Pat1a paralogue only found in vertebrate which localizes poorly to PB and has a separate function from the more conserved Pat1b protein (Marnef and Standart 2010). Although the data presented here neither confirms nor refutes the hypothesis, the substantial enrichment of PAT1 mRNA suggests that this transcript itself could play a role in PB assembly, function, or stability through a structural or catalytic activity.

The next most highly enriched RNA is the *RPM1* transcript, which is the catalytic RNA component of mitochondrial RNase P (Sulo *et al.* 1995). Although this RNA-protein complex has a well characterized mitochondrial function, we detected both *RPM1* and the protein component of the mitochondrial RNase P, Rpm2, in four of our six Dhh1-GFP IP samples. These results are also consistent with previous observations that the Rpm2 protein localizes to PB and genetically interacts with PB components (Stribinskis and Ramos 2007). Among the non-coding elements, two distinct classes of mitochondrially encoded ribozymes are also identified and confirmed by qRT-PCR (Figure S4), including several self-splicing introns from the *COX1* and 21S rRNA loci (*AI3, AI4, AI5*?, and *SCEI*). Only the DEAD-box helicase Mss116 that facilitates *in vivo* splicing of mitochondrial introns (Huang *et al.* 2005; Solem *et al.* 2006) is detected among the RBP co-isolating with Dhh1-GFP, while another mitochondrial helicase Suv3, involved in mitochondrial RNA decay, is not (Borowski *et al.* 2010; Bruni *et al.* 2012; Szczesny *et al.* 2013). Thus, PB/SG foci are enriched for a set of well characterized non-coding RNAs with catalytic activity along with their associated RBPs.

## DISCUSSION

Here, we have used a high-affinity antibody to isolate core Dhh1-GFP complexes at maximal yield under conditions that preserve the PB aggregate state. The method has allowed us to reproducibly isolate PB core components at level stoichiometric to that of the bait Dhh1-GFP as well as a number of PB/SG accessory proteins. Our results provide biochemical evidence for a number of proteins that have only been associated partially with PB/SG foci under certain conditions, or implicated in controlling PB/SG assembly by genetic studies without localization data (Buchan *et al.* 2013; Yang *et al.* 2014). SG proteins also co-purify with Dhh1-GFP, but at lower abundance than known PB proteins. Our results support the model of a continuum of mRNP granules transitioning between PB and SG foci, particularly during stress (Grousl *et al.* 2009; Buchan and Parker 2009). We expect that more proteins from our list might be identified to be functionally associated with Dhh1 in other non-stress condition screens (*e.g.* overexpressing or deletion mutants), since many proteins that regulate PB/SG function do not have well characterized RNA metabolism or translational regulatory activities (Buchan *et al.* 2013; Yang *et al.* 2014).

### LC-containing proteins and role of chaperones in regulating PB aggregation

PB formation is a dynamic stat between mRNP granule assembly and disassembly states during a stress response (Buchan 2014). In mammalian cells, a high proportion of RBP found in RNP granules contain LC domains that are necessary and sufficient to transform RNP complexes from soluble to aggregate states (Kato *et al.* 2012). Various heat shock proteins localize to SG and are required for disassembly and the re-initiation of translation upon removal from stress (Cherkasov *et al.* 2013; Buchan 2014). In parallel, prion protein assembly studies show that factors controlling protein aggregation depend on protein concentration, an organizing scaffold, and a network of chaperones (Summers *et al.* 2009). In a similar trend, 17% of the proteins in our PB enrichment are predicted to have LC/prion domains with more than two thirds being RBP. Furthermore, subgroups of protein chaperones are selectively enriched with Dhh1-GFP in the PB inducing condition of low glucose. Here we have shown that the Hsp40 chaperone Ydj1, which can bind to prion domains and also has a predicted LC domain, is specifically required for Dhh1-GFP foci assembly. Ydj1 is part of the Hsp40-70-110 chaperone network that regulates the aggregation-disaggregation of yeast prion proteins (Summers *et al.* 2009). The fact that Ydj1 affects PB foci formation strongly supports the model that Ydj1 mediates LC/prion domain interactions among RNP proteins to assemble PB. The accumulation of unfolded proteins during stress could affect the level of various chaperones in cells and drive the equilibrium toward aggregation of LC proteins into PB/SG (Buchan and Parker 2009; Buchan 2014). This model could explain why overexpression of Ydj1 alone does not cause Dhh1-GFP foci formation in the absence of an acute glucose stress. Ydj1 might affect PB assembly only by coordinating its activity with other co-chaperones, and without a stress trigger, the chaperone network equilibrium is not perturbed. Since Dhh1-GFP foci are still observed in some *ydj1*Δ cells grown to stationary phase, and Edc3-GFP foci are diminished but not completely missing in *ydj1*Δ cells exposed to glucose stress, there are likely multiple factors controlling PB/SG assembly.

### PB/SG and protein foci formation in response to stress

An important clue for understanding the relevance of Dhh1-GFP associated proteins with PB functions is the strong overlap with proteins that redistribute to cytoplasmic foci in response to various environmental stress (Narayanaswamy *et al.* 2009; Tkach *et al.* 2012; Shah *et al.* 2014). Transient protein aggregation is a physiological means to triage proteins for refolding or degradation (Escusa-Toret *et al.* 2013; Sontag *et al.* 2014). Major protein subgroups that are associated more with Dhh1-GFP in the PB induced state (*e.g.* chaperones, metabolic enzymes involved in amino acid/purine synthesis, tRNA synthetases) also form foci in stationary phase cells. The majority of these proteins form aggregates that are reversible, suggesting that they are not permanently denatured protein aggregates (Narayanaswamy *et al.* 2009). Protein granules observed in stationary phase cells are associated with a network of chaperones that are very similar to those we see associated with Dhh1-GFP (O’Connell *et al.* 2014). Cells might assemble metabolic enzymes into foci to modulate their activities, to provide structural stability, or to concentrate metabolites and RNA into transient storage in order to quickly re-enter the cell cycle once stress is removed (O’Connell *et al.* 2012). Most of the proteins in our isolations that form foci also bind ATP (*i.e.* metabolic enzymes and tRNA synthetases), and almost half also bind RNA. According to the REM hypothesis (Scherrer *et al.* 2010; Hentze and Preiss 2010), metabolic enzymes that bind to mRNA as well as metabolites such as ATP can provide cells a mechanism to link post-transcriptional regulation with cellular metabolism. Cells might be able to enhance long-term survival by allowing metabolic enzymes the ability to quickly adapt to binding mRNA when substrate availability is limited during starvation. Finally, a subgroup of tRNA synthetases is co-precipitated with the foci-forming tRNA synthetase Ils1-GFP in stationary phase cells (O’Connell *et al.* 2014) very similarly with the tRNA synthetase subgroup that interacts with Dhh1-GFP in acute glucose depletion. Regulation of the nuclear tRNA pool has been shown to be coordinated with PB assembly in response to amino acid starvation in yeast (Hurto and Hopper 2011). The tRNA synthetase chaperone Arc1 is also coisolated with Dhh1-GFP and is required for SG assembly (Yang *et al.* 2014). A current model suggests that cells modulate the aminoacyl-tRNA repetoire to regulate preferential translation of certain proteins during stress (Subramaniam *et al.* 2014) or during cell proliferative state (Gingold *et al.* 2014).

### RNA associated with Dhh1-GFP complexes

The reproducible isolation of subsets of RBP in our PB enrichment allows us to characterize the associated transcripts specific to Dhh1-GFP subcomplexes. The paucity of only 79 transcripts enriched is conceivably due to the complexity of the RNA mixture that is coisolated with Dhh1-GFP. It is estimated that 70% of cellular mRNA in yeast is associated with polyribosomes (Arava *et al.* 2003), and since ribosomal subunits are co-isolated with Dhh1-GFP along with RNP complexes, the cellular mRNAs associated with ribosomal subunits can increase the overall background of RNA enrichment and decrease the effective enrichment of RNAs associated with RNP complexes. However, as the majority of RNA coisolating with Dhh1-GFP are noncoding, the identified transcripts are not isolated simply because of ribosome association.

We also show for the first time a physical association between PB and self-splicing introns within the mitochondrial *COX1* and 21S rRNA loci. Consistent with these findings, PB has been implicated in the splicing of mitochondrial introns as the respiratory deficiency of *dhh1*Δ and *lsm6*Δ mutants are rescued by deletion of the self-splicing mitochondrial introns (Luban *et al.* 2005). Another relevant aspect of self-splicing introns association with PB is their ability to act as mobile elements within the mitochondrial genome (Moran *et al.* 1992; Yang *et al.* 1996; Nielsen and Johansen 2009). PB components are required for retrotransposition of nuclear Ty elements that also localize to PB (Checkley *et al.* 2010). It is plausible that PB plays a role in regulating the mobility of self-splicing introns within the mitochondrial genome.

In conclusion, we have identified a class of proteins that controls PB formation, other classes that link PB assembly to other stress response protein foci, as well as several catalytic RNP complexes that connect PB with mitochondrial RNA processing. By isolating and determining the composition of PB in differential assembly conditions, we have identified components that would be undetectable by microscopy either because of low abundance, transient interactions, or association in non-stress conditions when foci are undetectable. These results demonstrate the usefulness of a global biochemical approach, complementary to cytological and genetic studies, which can enhance the understanding of RNP granule assembly and function.

## ACKNOWLEDGMENTS

We thank Cecilia Garmendia-Torres for providing strains; John Aitchison for use of the planetary ball-mill grinder; Mark Gillespie for advice on antibody crosslinking; Gareth Cromie, Bruz Marzolf, and Pamela Troisch for advice on microarray analyses; Young-Ah Goo, David Goodlett, Deborah Chang, and Ryan Austin for advice on mass spectrometry analyses; Claire Gustafson for assistance in the early stages of the project; Cecilia Garmendia-Torres, David Morris, David Goodlett, John Aitchison, Dan Jarosz, and members of the Dudley lab for helpful discussions; and Cecilia Garmendia-Torres and John Woolford for helpful comments on the manuscript. Mass spectrometry was performed by Young-Ah Goo at the University of Washington School of Pharmacy’s Mass Spectrometry Facility.

